# Analyzing the symmetrical arrangement of structural repeats in proteins with CE-Symm

**DOI:** 10.1101/297960

**Authors:** Spencer E Bliven, Aleixa Lafita, Peter W Rose, Guido Capitani, Andreas Prlić, Philip E Bourne

## Abstract

Many proteins fold into highly regular and repetitive three dimensional structures. The analysis of structural patterns and repeated elements is fundamental to understand protein function and evolution. We present recent improvements to the `CE-Symm` tool for systematically detecting and analyzing the internal symmetry and structural repeats in proteins. In addition to the accurate detection of internal symmetry, the tool is now capable of i) reporting the type of symmetry, ii) identifying the smallest repeating unit, iii) describing the arrangement of repeats with transformation operations and symmetry axes, and iv) comparing the similarity of all the internal repeats at the residue level. `CE-Symm 2.0` helps the user investigate proteins with a robust and intuitive sequence-to-structure analysis, with many applications in protein classification, functional annotation and evolutionary studies. We describe the algorithmic extensions of the method and demonstrate its applications to the study of interesting cases of protein evolution.

**Availabilit**: `CE-Symm` is an open source tool integrated into the BioJava library(www.biojava.org)and freely available at https://github.com/rcsb/symmetry.

## Author summary

Many protein structures show a great deal of regularity. Even within single polypeptide chains, about 25% of proteins contain self-similar repeating structures, which can be organized in ring-like symmetric arrangements or linear open repeats. The repeats are often related, and thus comparing the sequence and structure of repeats can give an idea as to the early evolutionary history of a protein family. Additionally, the conservation and divergence of repeats can lead to insights about the function of the proteins.

This work describes `CE-Symm` 2.0, a tool for the analysis of protein symmetry. The method automatically detects internal symmetry in protein structures and produces a multiple alignment of structural repeats. The algorithm is able to detect the geometric relationships between the repeats, including cyclic, dihedral, and polyhedral symmetries, translational repeats, and cases where multiple symmetry operators are applicable in a hierarchical manner. These complex relationships can then be visualized in a graphical interface as a complete structure, as a superposition of repeats, or as a multiple alignment of the protein sequence. `CE-Symm` 2.0 can be systematically used for the automatic detection of internal symmetry in protein structures, or as an interactive tool for the analysis of structural repeats.

## Introduction

François Jacob described molecular evolution as a “tinkering” process, where pre-existing elements are combined and repurposed to solve new biological problems [1]. Traces of this “tinkerer evolution” can be seen in the widespread reuse of structural elements in proteins at different scales: small motifs [2], functional domains [3], and protein oligomerization [4]. One example is the repetition of structural elements within a protein chain, thought to arise from gene duplication and fusion events [5].

It is common for structural repeats in proteins to maintain a symmetric arrangement [6], which has been associated with many biological functions [7]. The internal symmetry of proteins is thought to arise from ancestral quaternary structures fused into a single polypeptide chain [8–10]. However, since symmetric protein folds theoretically have a folding thermodynamic advantage, their symmetry could also have arisen by evolutionary convergence [11]. On the other hand, the evolution of functional patches is often symmetry breaking [12]. High-quality alignments of structural repeats are essential to resolve these opposing evolutionary explanations and understand the tension between conservation and divergence.

There are a number of computational methods and tools to detect and analyze structural repeats in proteins. Some methods focus on the detection of patterns and periodicities in protein structures and are better suited for the prediction of solenoids repeats [13–16]. Other methods, including the `CE-Symm` tool presented here, make use of structural alignments and generally perform better in larger regular repeats [6, 17–21]. These two approaches have also been combined to improve the repeat detection performance [22]. In addition, there are tools that use existing libraries of protein structural repeats [23,24]. The most comprehensive database of known structural repeats is `RepeatsDB` [25].

Two of the repeat detection methods primarily focus on the detection of internal symmetry. Both `SymD` [20] and `CE-Symm` [21] start with a self-alignment of the structure against itself to identify significant self-similarities. The extraction of repeats from the self-alignments, however, is a nontrivial task, so initial versions of both methods were concerned only with internal symmetry detection (binary decision) and estimation of the number of repeats. Here we present an extension of `CE-Symm` (version 2.0) that, apart from accurately detecting internal symmetry in proteins, defines the repeat boundaries, reports the type of symmetry and describes the arrangement of repeats using symmetry axes. The similarity of the structural repeats can be further compared at the residue level in a multiple structure alignment.

### Types of symmetry

Several definitions of **internal symmetry** and **repeats** are possible, depending on the biological question of interest. For the purposes of this paper, we define it as the regular arrangement of a common repeating structural unit within a protein chain. Therefore, a repeat is an asymmetric structural motif present multiple times in the same structure. We restrict our consideration of repeats to cases where the orientation between adjacent structural units is regular; that is, where a consistent geometric transformation can be applied to superimpose each repeat onto the next. In other words, `CE-Symm` focuses on identifying repeats which conserve both the structure and the interface between repeats.

Several types of internal symmetry can be derived from this broad definition. The most basic division is between **closed symmetry** and open symmetry. In proteins with closed symmetry, the repeats are arranged in a point group symmetry. This can be defined mathematically as a set of rotations that superimpose equivalent repeats while keeping at least one point at the center of rotation fixed. In contrast, repeats in proteins with open **symmetry** are related by transformations with a translational component. Examples of closed and open symmetry can be found in Fig 1a-d and Fig 1e-h, respectively.

**Fig. 1.**
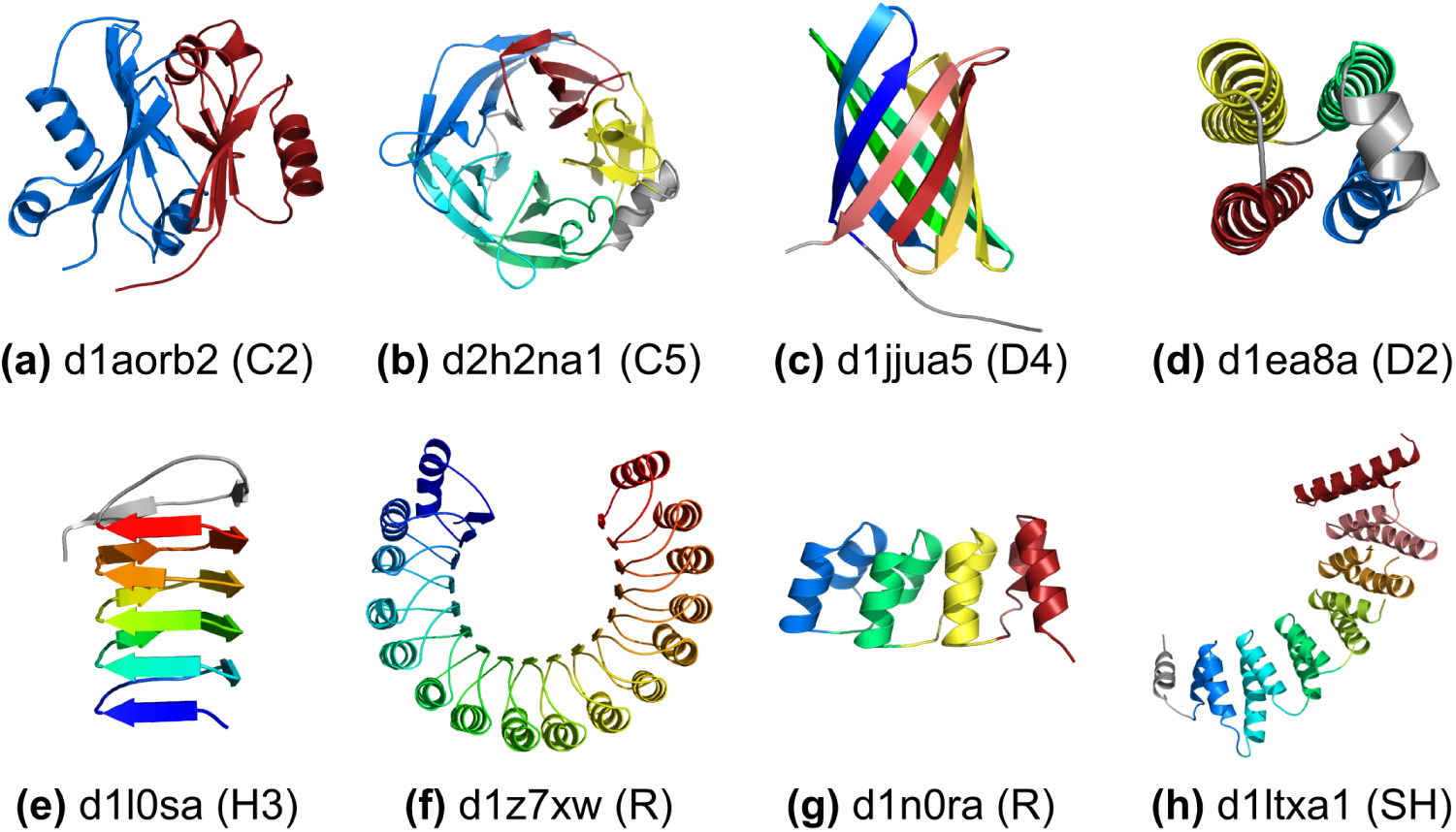
Examples of protein globular domains with internal symmetry. Protein domains are labeled with SCOP domain identifier [26]. a) N-terminal domain of aldehyde ferredoxin oxidoreductase with 2-fold rotational symmetry (C2) of an alpha+beta motif; b) 5-bladed beta propeller with a helical insertion between second and third blades with 5-fold rotational symmetry (C5) of a 4-stranded beta-sheet motif; c) Beta-barrel with 8 beta strands in a 4-fold dihedral symmetry (D4) of single stranded motifs; d) 4-helical bundle with dihedral symmetry connectivity (D2) of single helical motifs; e) Beta-helix with single stranded right-handed helical symmetry (H3); f) Leucine rich repeats with open rotational symmetry (R) of 16 up and down alpha-beta motifs; g) Designed Ankyrin repeat protein with 4 translational repeats (R) of double helical motifs; h) Alpha-alpha right-handed superhelix (SH) of double helical motifs. Repeats are colored from blue, N-terminal, to red, C-terminal. Non-repeating parts of the structure are colored in grey.

Closed symmetries can be further characterized according to the possible chiral point groups: cyclic (C*n*), generated by a single n-fold rotational operator (Fig 1a-b); dihedral (D*n*), which requires an *n*-fold rotation and *n* perpendicular 2-fold operators (Fig 1c-d); and polyhedral point-groups (T, O, and I), which feature non-perpendicular rotation operators. Both cyclic and dihedral internal symmetries are common in proteins, but, although common at the quaternary structure level, polyhedral symmetries have not yet been observed within a single polypeptide chain.

Open symmetry can be further subdivided into special cases of helical, translational, and superhelical repeats. Helical symmetry consists of repeats arranged around a screw axis, where each repeat is related to the next by a fixed linear translation combined by a rotation around the central axis (Fig 1e). In cases where the rotation angle is close to an fraction of a turn, we indicate the approximate number of subunits needed per turn (H*n*). Proteins with open symmetry that have negligible translation are called rotational repeats (Fig 1f), and those with negligible rotation between repeats are called translational repeats (Fig 1g), both annotated as R. Superhelical symmetry (SH) provides the most general description of repeats with open symmetry, and is reserved for cases which cannot be expressed as a single fixed operator relating each repeat to the next. Instead, the rotation axis between adjacent repeats precesses along a helical path (Fig 1h). Proteins with open symmetry are sometimes referred to as solenoid proteins [27].

## Methods

`CE-Symm` analyzes the symmetry in a protein structure and produces a multiple alignment of all repeats, as well as ancillary information about the type and order of symmetry in the structure. An overview of `CE-Symm` 2.0 alignment steps is shown in Fig 2. These are described in detail below, but consist of (1) structural self-alignment, (2) order detection, (3) refinement to a multiple alignment, (4) Monte Carlo optimization of the multiple alignment, and (5) point group symmetry detection. These steps are repeated iteratively to detect multiple levels of symmetry (hierarchical symmetry) and higher-order point groups.

**Fig. 2.**
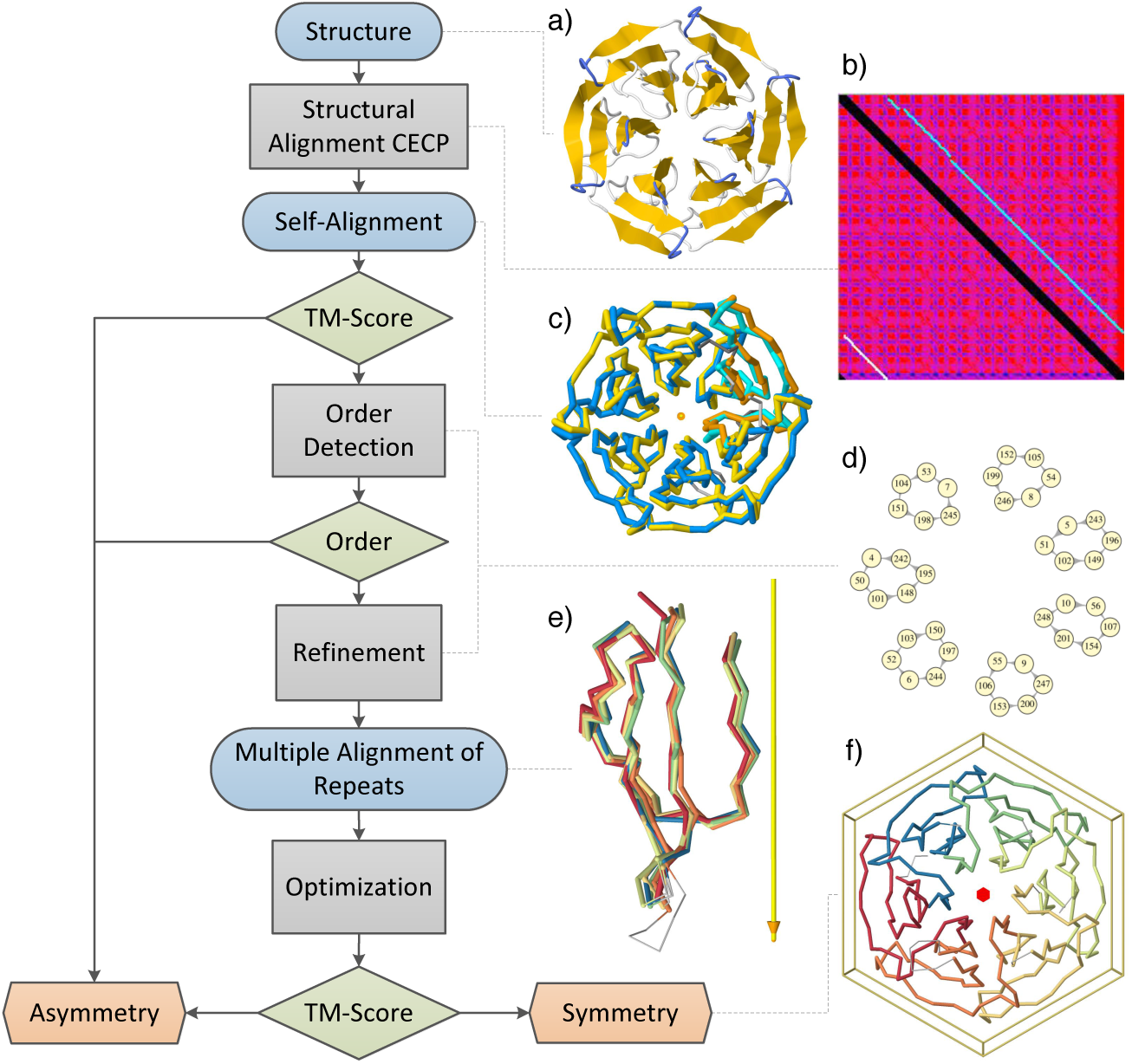
Flowchart of one iteration of the CE-Symm algorithm. Algorithm steps are grey rectangles, inputs and outputs are blue rounded rectangles, decision rules are green rhomboid boxes and final classifications are orange hexagonal rectangles. Additional iterations on the resulting repeats may be performed to detect further symmetry axes or hierarchical symmetry. The images on the right represent, from top to bottom: a) initial structure, colored by secondary structure elements; b) self-alignment dot-plot matrix, where similarity score is a range from blue (high similarity) to magenta (low similarity), the identity alignment is blacked out and the optimal self-alignment path is in white; c) superposition of the structure against itself based on the optimal self-alignment, where the original structure is in blue and cyan and a copy of the structure is in yellow and orange (orange and cyan correspond to the regions of the alignment involving a circular permutation); d) subset of the alignment graph with seven connected components of six aligned residues each; e) superposition of the six internally symmetric repeats according to the symmetry axis (yellow bar) and their residue equivalencies; and f) structure inside the six-fold cyclic symmetry (C6) box, with repeats colored differently.

### Self-alignment

`CE-Symm` begins with a structural self-alignment (other than the identity alignment) of the input protein structure using the Combinatorial Extension (CE) algorithm [28] (Fig 2b-c). Identifying significant self-alignments was the primary focus of the first version of the algorithm [21]. In the self-alignment of structures with closed symmetry the first and last repeats are aligned, forming a circular permutation (CP) of the structure. This is why the structure alignment method used in `CE-Symm` shares algorithmic primitives with `CE-CP` [29]. For proteins with open symmetry, the initial self-alignment will always be missing one of the repeats due to the translation component of the symmetry operator.

The alignment quality is quantified using TM-Score [30]. Both irregularly arranged repeats and large asymmetric regions in a structure will reduce the score of the self-alignment. In addition, open symmetry will generally have lower scores than closed symmetry, because the terminal repeats are unaligned in the initial self-alignment.

### Order of symmetry detection

The order of symmetry is defined as the number of symmetric units (repeats) in a structure. Extracting the order of symmetry is a key part of symmetry detection, and subsequent steps of the `CE-Symm` method depend on its correctness.

Two methods to automatically determine the order of symmetry in closed structures were described in the previous `CE-Symm` publication [21]:DeltaPositions and RotationAngle. An error in the distance formula was corrected in `CE-Symm` 2.0 (see S1 Text, Supplemental Methods), but the DeltaPosition method still gives better overall performance and is used by default for closed symmetry.

A third method for order detection which is able to handle open symmetry has been introduced in the new version of the tool and is named GraphComponent. Conceptually, the self-alignment is treated as a directed graph over the set of aligned residues (Fig 2d). Residues that are aligned in all *k* repeats will form a path with *k* nodes. For open symmetry these paths tend to be disjoint, so simply finding the most frequent size of the connected components in the graph can accurately determine the order for open symmetry. For well-aligned cases of closed symmetry, the aligned residues form a cycle of *k* nodes, so the same method can also work in the general case. Those residues which participate in a path or cycle of the most frequent size form the refined alignment discussed in the following section.

For cases of closed symmetry, small alignment errors can lead to a situation where paths of *k* residues do not form a closed cycle, but rather lead to a different residue at a small offset in the sequence. This can lead to failures of the GraphComponent order detector due to the merging of multiple alignment paths. This case can be handled by the DeltaPosition order detector.

Whether a protein is open or closed can be easily determined by looking for a circular permutation in the self-alignment. `CE-Symm` uses DeltaPosition for closed cases where a permutation is found and GraphComponent for open cases. This can also be overridden by the user when running `CE-Symm`.

### Refinement to a multiple alignment

The refinement procedure takes as input the self-alignment of the structure and the order of symmetry (*k*) and returns a multiple alignment of the repeats (Fig 2e). CE-Symm has two implementations of the refinement procedure: GraphComponent and DeltaPosition, which are closely related to their respective order detectors.

The GraphComponent refiner combines all connected components of the self-alignment graph with size equal to the order of symmetry. Each connected component contributes one column to the refined alignment of repeats. Care must be taken that the repeat sequences preserve the sequence order of the polypeptide chain; where some pairs of connected components would violate this property, some are discarded in a way that maximizes the total length of the resulting alignment.

The DeltaPosition refiner takes the self-alignment graph and modifies it until all remaining nodes are part of *k*-cycles. The modification heuristic is described in detail in the Supplemental Methods (S1 Text). These cycles each contribute one column of the multiple alignment of the symmetric repeats, as in the GraphComponent refiner. Note that, like the GraphComponent refiner, the multiple alignments obtained at the end of this stage consist of ungapped columns, so all repeats are of the same size.

### Optimization

The multiple alignment obtained from the refinement is sometimes far from optimal, and depends very much on the quality of the self-alignment. In addition, the refinement process prioritizes precision over coverage, which means that only the best residue equivalencies will be included, resulting in a shorter multiple alignment. The goal of the optimization is to increase the multiple alignment length while keeping the RMSD low (Fig 2f). Furthermore, the optimization procedure can improve parts of the alignment that were not fully represented in the self-alignment, and thus not captured in the refinement result.

The optimization process uses a similar approach to the Combinatorial Extension Monte Carlo (CEMC) multiple structure alignment algorithm [31]. The multiple alignment can be described by a matrix, where the rows represent aligned structures and the columns represent aligned positions (residue equivalencies). Rearranging and modifying the entries of the matrix results in changes of the multiple alignment. There are four possible moves (changes in the multiple alignment):

1. **Expand:** increase the alignment length by extending the boundary of a group of sequential residues, chosen randomly. This move requires the addition of an alignment column.
2. **Shrink:** decrease the alignment length by decreasing the boundary of a group of sequential residues, chosen randomly. This move requires a deletion of an alignment column.
3. **Shift:** move a group of sequential residues, chosen randomly, of one structure (row), chosen randomly, one position to the right or to the left.
4. **Insert gap:** delete one entry of the matrix, chosen randomly. This is equivalent to inserting a gap in one residue position (column) of one structure (row).

The insertion of gaps allows for partial repeat similarities in the alignment. All moves take into consideration that rows of the alignment occur sequentially in the protein sequence, so unaligned residues between repeats can be considered either at the end of a repeat or the beginning of the following one. In addition, the shrink and insert gap moves have been biased, so that the probability of choosing an alignment column or an equivalent residue, respectively, is proportional to the average inter-residue distance of the given column or the given residue, respectively. A geometric distribution with parameter 0.5 is chosen to allocate the probability among alignment columns. A schematic representation of the steps and how they affect the multiple alignment is provided in S1 Fig.

After each optimization step, an alignment score is calculated. The score function to be optimized has also been smoothed with respect to the original CEMC score to remove discontinuities:

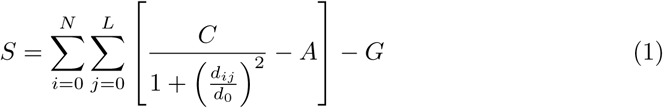

*N* is the number of structures (rows) in the alignment; *L* is the number of equivalent positions (columns) in the alignment, including gaps; *C* is the maximum score of an alignment position (by default set to 20);*d*_*ij*_ is the average distance from aligned residue *j* in structure *i* to all its equivalent residues; *d*_0_ is the structural similarity function parameter, as defined by the TM-score [30]; A is the distance cutoff penalization, which shifts the function to negative values when the maximum allowed average distance of an aligned position (*d*_*c*_) is higher than *d*_*ij*_; and *G* is a linear gap penalty term. Calculation of *A* using a distance cutoff parameter *d*_*c*_ (by default set to 7Å) is straightforward from the condition that the score *S*d has to be 0 when *d*_*ij*_ = *d*_*c*_. The shape of the score function for different values of *d*_*c*_ is shown in S2 Fig.

Moves with positive score changes are always accepted. The acceptance probability of a negative scoring move is proportional to the score difference and decreases proportional to the number of optimization steps as follows:

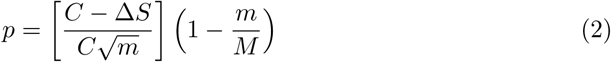

*m* is the current iteration number, ΔS is the change in alignment score and *M* is the maximum number of iterations. The maximum number of optimization iterations is proportional to the length of the protein, by default a hundred times the number of residues in the protein. Optimization finishes either because it reaches the maximum number of iterations or in the case that no moves are accepted for a fraction of the total number of iterations (by default *M* divided by 50).

### Recursive symmetry detection

So far, the procedure described can only identify symmetry operations that require a single axis. However, some structures present symmetries represented with more than one axis. This is the case for point groups other than cyclic, like dihedral symmetry, or structures with more than one level of symmetry, what we define as hierarchical symmetries. Multiple CE-Symm iterations are run in a recursive manner, i.e. repeats found in previous rounds are recursively fed into the next run until a non-significant result (no symmetry) is found. The goal is to find all the significant symmetry levels of a structure.

At the end of an iteration, repeats are extracted from the internal symmetry result and one of them is chosen as the representative, by default the N-terminal repeat. Results of successive iterations are merged by combining the symmetry axes and multiple alignments, generating a unique result for the query structure.

The recursive symmetry detection allows better order determination for difficult cases (e.g., TIM barrels), because usually fractions of the order of symmetry are initially found (e.g., 2-fold instead of 8-fold). Continuing the analysis recursively breaks the structure down to the true asymmetric repeating units (e.g., with three levels of symmetry: 2-fold, 4-fold and finally 8-fold).

### Significance

There are three decision checkpoints in the algorithm flowchart in Fig 2. The first significance criterion for a symmetry result is the self-alignment TM-score. Like in the previous version, the default threshold value is set to 0.4. The second significance criterion is the order of symmetry. A symmetric structure must have symmetry order greater than one and the refinement of the self-alignment into a multiple repeat alignment has to be successful. The third significance criterion is the average TM-score of the multiple alignment of repeats, defined as the average TM-score of all pairwise repeat alignments. The default threshold value for the average TM-score is set to 0.36, because a 10% decrease from the original TM-score is allowed after refinement due to the restrictive conditions imposed on it. In addition the number secondary structure elements (SSE) of the final asymmetric repeating unit is considered. If the the number of SSE of each repeat is lower than the threshold, the result will be considered non-significant. For many applications it may be desirable to exclude simple repeat units (e.g. helical bundle proteins), but these are included in CE-Symm analysis by default in order to find the highest possible symmetry in a structure.

### Symmetry type determination

The recursive symmetry detection identifies a collection of symmetry axes that describe the arrangement of repeats in the query structure. In many cases, several of these axes can be combined to form higher-order symmetries. For example, a two-fold rotation axis can be combined with another orthogonal axis to form dihedral symmetry. Near-identical rotation axes can also be combined to form higher-order rotational symmetry.

To determine the point group symmetry, we build on the algorithm described by Levy *et al*. [32]. The symmetry axes can be found efficiently by first considering only the centroids of each repeat, since they must be in a symmetric configuration if the entire complex is symmetric. To find all possible symmetry axes, the centroids are rotated around axes that go through the centroid of the whole structure using an orientation grid search in quaternion space [33]. For each orientation, the RMSD of the aligned centroids is calculated. If the centroids align within a threshold, then all C*α* atoms are superimposed. The symmetry axis is then defined by the rotation matrix of this superposition. If the RMSD is less than a threshold value (i.e., 5Å), the symmetry operation is considered valid. Since symmetry operations form a group, only a few are needed to complete the full point group

This procedure allows the combination of axes that have been considered separately by `CE-Symm`. The point group is included in the final symmetry output and displayed to the user as a polyhedron box around the protein structure.

### Benchmarking datasets

For evaluation purposes we used the manually curated dataset of 1,007 domains selected randomly from the set of SCOP superfamilies, introduced in our previous study [21]. The benchmark is intended to be representative of structural domains in the PDB, and repeats are present in 25.8% of domains. Domains were curated to require reasonably high coverage by repeats with conserved topology, but allowing for structural divergence and flexibility. A small number of classifications were updated to be more consistent with the new symmetry definitions, especially for the cases of open symmetry. The updated version of the internal symmetry dataset (v2.0), together with the reasons of the modified annotations, is summarized in S1 Tab. The benchmark dataset and results are available in S2 Tab or https://github.com/rcsb/symmetry-benchmark. An important note is that in the evaluation of the previous version the open symmetry cases in the benchmark were part of the asymmetric (negative) set, while they are part of the symmetric (positive) set in this evaluation.

`RepeatsDB` was used as an additional benchmark of positive cases [25]. At time of download, `RepeatsDB` contained 3,689 manually reviewed entries (accessed October 18, 2018). Entries consist of single chains, and may contain multiple structural domains or multiple repeat regions. We selected a total of 3,503 chains with repeats of classes III (solenoid repeats) and IV (closed repeats) that were part of the `RepeatsDB-lite` benchmarking dataset [24]. Chains which were either annotated in RepeatsDB as having multiple repeat regions or in ECOD [34] as having multiple structural domains were considered multidomain chains in the analysis. The `RepeatsDB` benchmarking dataset and `CE-Symm` results are listed in S3 Tab.

## Results

### Method evaluation

In our previous article we compared the performance of CE-Symm against SymD. The performance in symmetry detection has only been affected by the additional order detection and alignment refinement steps. The ROC curves of both versions are very similar, with a slight reduction of false positives in the new one (S3 Fig). At the default TM-score threshold values for result significance, the false positive (FP) rate has decreased from 5.5% to 2.5%, while the true positive (TP) rate has been reduced from 81% to 76% on the benchmarking dataset. The bottleneck in symmetry detection continues to be finding a significant self-alignment.

The different methods for order detection perform similarly for closed symmetry cases in the benchmark (S4 Tab). The simpler GraphComponent method performs worse than the others, but it is the only one that can be used for open symmetries, while the DeltaPosition detector performs better than the RotationAngle method, particularly for difficult cases.

On average for symmetric entries in the benchmark where `CE-Symm` could find symmetry, the optimization step extended the repeat length by 43%, reduced the RMSD by 1.8% and increased the average TM-Score of the repeat alignment by 19.6%. Furthermore, using optimization an additional 23 cases (9% of the symmetric structures in the benchmark) were correctly identified as symmetric (209 with optimization, 186 without), which is a 12% improvement in symmetry detection. Because the highest scoring alignment of the simulation trajectory is taken as the result, optimization can only improve the initial alignment.

A detailed evaluation of predicted number of repeats by `CE-Symm` is shown in S3 Fig. Among incorrect predictions, `CE-Symm` tends to underpredict the number of repeats, typically as a fraction of the true number of repeats (e.g. predicting 4 repeats for a C8 TIM-barrel). Examples with high-order rotational symmetry are also often missed due to the default maximum order of 8 used in the DeltaPosition method. Overall, `CE-Symm` 2.0 is able to predict the number of repeats correctly in 89% of cases. High order open symmetries remain challenging due to the prevalence of kinks and structural inhomogeneities in large open structures.

The `RepeatsDB-lite` method was also run on the benchmark for comparison (S5 Fig). `RepeatsDB-lite` only detects proteins with three or more repeats, so one and two-repeat cases in the benchmark were binned together for analysis. With this definition, it predicts the correct number of structures for 72% of the benchmark. Since the method is based on a library of known repeat units, it tends to miss repeats in benchmark cases that are not similar to the training dataset. Additionally, the method does not enforce high coverage or symmetric orientations between repeats. This is desirable when identifying candidates that may have repeats, but it leads to a rather high false positive rate (24%) relative to the definitions of repeats used when curating the benchmark. Raw data for the benchmark results is available in S3 Tab and at https://github.com/rcsb/symmetry-benchmark.

`CE-Symm` was run on the `RepeatsDB` dataset to assess its performance on a larger set of symmetric proteins. It was able to detect repeats in 69% of the dataset of `RepeatsDB` reviewed entries. Considering only single-domain proteins improves the recall of 77%, indicating that multi-domain proteins are challenging for the method. The database classifies repeat regions according to the Kajava tandem repeat classes [35]. `CE-Symm` achieves a recall of 64% for single-domain solenoid repeats (class III) and 89% for closed repeats (class IV). Thus, `CE-Symm` performs best for closed repeats with single-domain input.

### Sequence-structure analysis

The new `CE-Symm` tool is capable of presenting internal symmetry as a multiple structural alignment of repeats, which enables direct association of sequence and structure and can be used for comparative and evolutionary analyses of protein structures. The superposition used for the alignment is constrained by the axes of symmetry found in the structure, so that equivalent residue positions maintain the symmetric orientation. Symmetry-aware alignments are important to study, for example, binding and active sites at the internal symmetry interface, like calcium binding in *βγ*-crystallins (Fig 3).

**Fig. 3.**
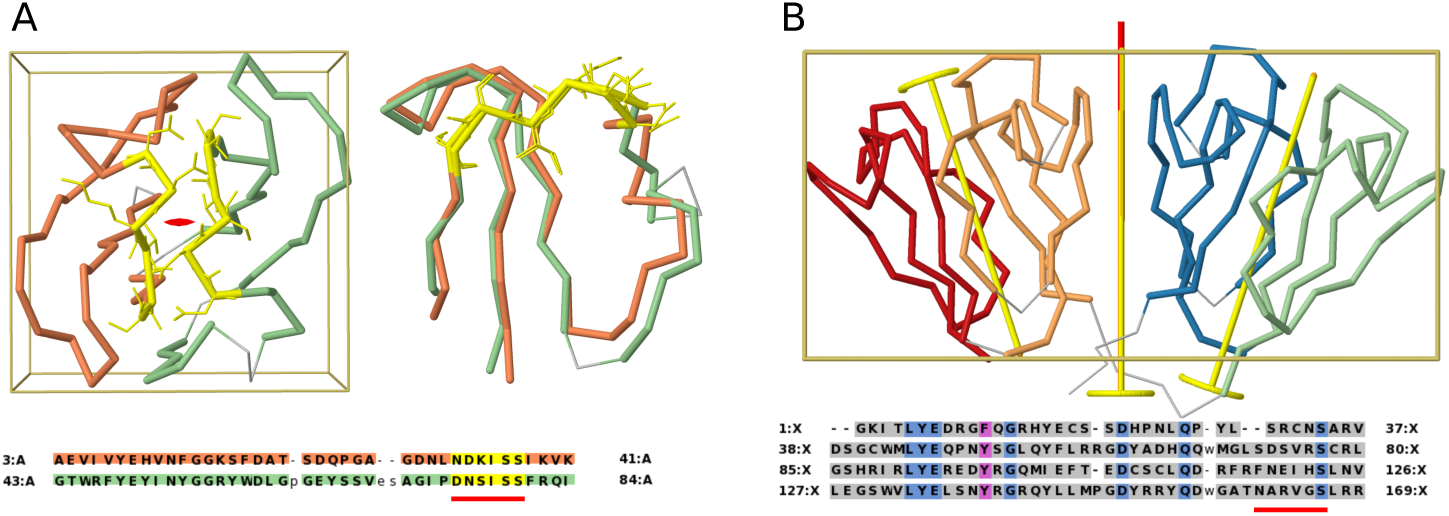
Internal symmetry in crystallin proteins. A) An archaean *βγ*-crystallin with two repeats per chain (3HZ2). The full chain is displayed along its 2-fold axis, followed by a superposition of the repeats. The conserved calcium binding motif N/D-N/D-#-I-S/T-S is highlighted in yellow throughout. B) Human *γ*-D crystallin structure with four repeats per chain (1HK0). Two levels of symmetry exist: a C2 symmetry within each globular domain, and an additional C2 axis relating the two domains. The calcium binding motif has been lost (red bar below sequence), but other conserved positions (blue and magenta in the sequence) show the homology between the repeats.

Structural alignment of the repeats can reveal conserved motifs that have persisted since the duplication event. One example is the *βγ*-crystallin superfamily, which occurs in a variety of repeat arrangements. Many *βγ*-crystallins contain a calcium binding site motif [36]. As shown in Fig 3A, the calcium binding motif is structurally conserved after a 2-fold rotation around the symmetry axis, and the residue side-chains preserve their orientation. Furthermore, calcium coordinates residues from both repeats, making the two-fold symmetry an essential feature of the binding site.

On the other hand, duplication events allow the appearance of asymmetry by independent sequence and structural divergence of the repeats. An example is the MaoC-like thioesterase/thiol ester dehydrase-isomerase superfamily (SCOP: d.38.1.4). Members of this family fold into a characteristic ‘hot dog fold’ which binds coenzyme A and catalyzes the dehydration of various bound fatty acids. Typically the MaoC-like proteins contain one hot dog domain per chain and assemble into dimers, tetramers, or hexamers [37]. Some members of the family contain a duplication of the hot dog fold [38], accompanied by the loss of the catalytic motif R/N-####-H in one of the domains, in order to accommodate bulkier substrates which would otherwise not fit in a single domain [37]. The structural divergence of the catalytic site in one of the repeats of the double hot dog subunit can be easily observed with `CE-Symm` (Fig 4).

**Fig. 4.**
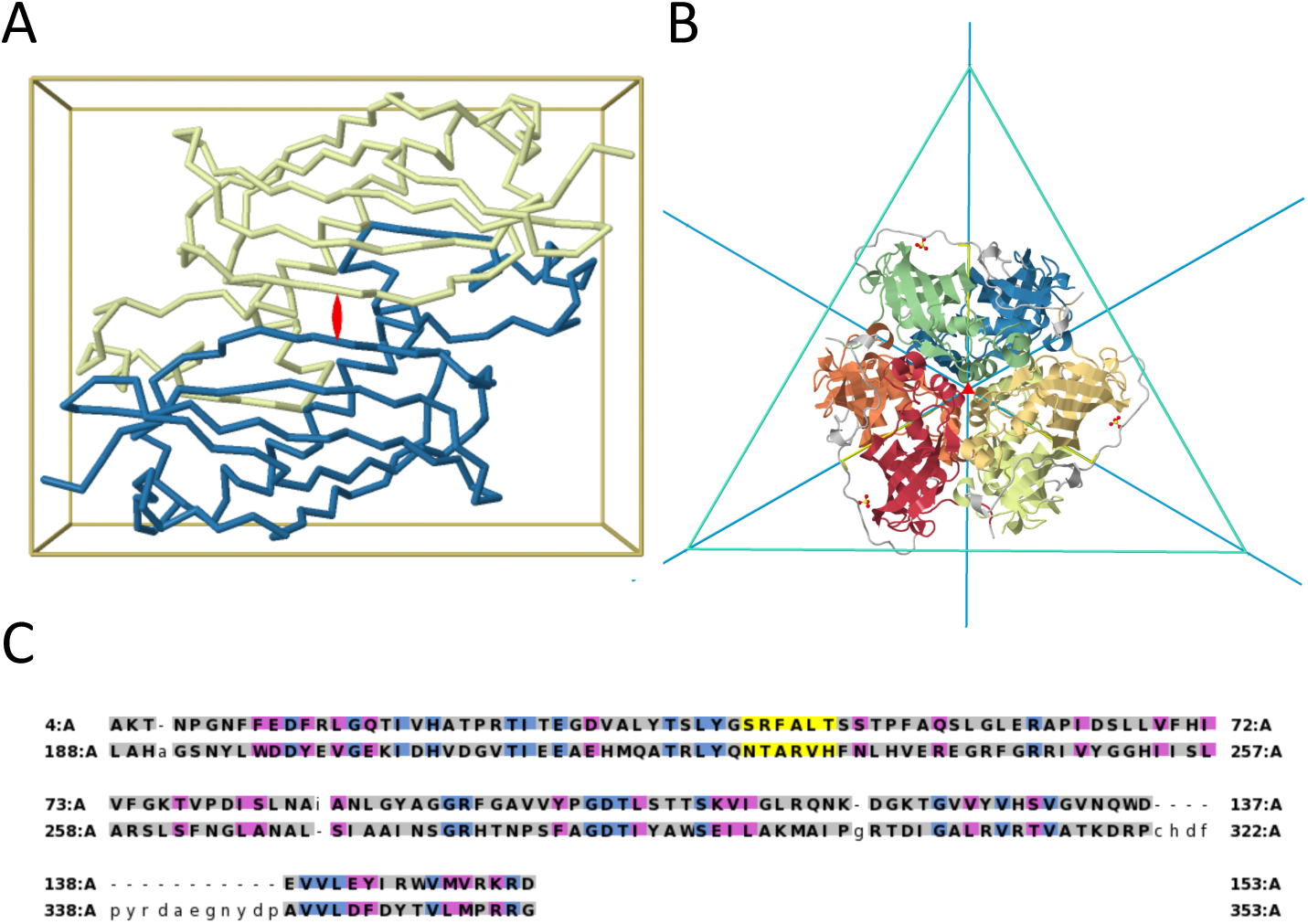
Hot dog fold duplication. A) Internal C2 symmetry in one chain of a MaoC domain protein dehydratase from Chloroflexus aurantiacus (4E3E) displaying a “double hot dog” fold. B) Full trimeric assembly, with the six individual hot dog domains colored. The quaternary structure has a threefold cyclic symmetry that combines with the twofold internal symmetry into a dihedral D3 symmetry equivalent to the quaternary structure of homologs without the internal domain duplication. C) Sequence alignment showing that the catalytic R/N-####-H motif (yellow) is lost in the first domain but retained in the second. Amino acid identity is shown in blue and similarity in magenta.

### Multiple levels of symmetry

Some proteins contain more than one axis of symmetry. In those cases, the axes of symmetry can be collinear, orthogonal or independent to each other. If the axes are collinear, they can be combined into a single axis with higher symmetry order. If the axes are orthogonal, they can be combined into a point group of higher symmetry order.

If the axes are independent to each other, multiple levels of symmetry exist in the structure in a hierarchical organization. This can be an indication of multiple independent duplication events, like in the case of *γ*-crystallins (Fig 3B), where four repeats are related by two independent 2-fold axes corresponding to two successive duplication events.

### Internal symmetry and assembly stoichiometry

Additionally, the internal symmetry axes can also combine with the quaternary symmetry axes. Therefore, internal symmetry can increase the order of symmetry of a protein complex. Returning to the previous MaoC-like protein example, the internal two-fold axis of the double hot dog domain in Fig 4B is orthogonal to the three-fold quaternary symmetry axis, combining for an overall dihedral symmetry. This arrangement is structurally similar to the D3 quaternary symmetry of hexameric single hot dog proteins (e.g. 1YLI). Accounting for internal symmetry when comparing the two protein assemblies is therefore important, because proteins can have a similar overall structure despite their different subunit compositions. It would be misleading to say that the structure of the trimeric and hexameric MaoC-like proteins are substantially different. Another well-know example of similar overall arrangement with different subunit composition are DNA clamps, which promote processivity in DNA replication. In archaea and eukaryotes, the clamp is a trimer, while in bacteria it is a dimer [39]. Furthermore, all DNA clamps have further internal symmetry axes leading to an overall D6 symmetry. As a historical note, the homology between bacterial and eurkaryotic DNA clamps was only acknowledged when the structures were solved and the similarity of their complexes was identified [40].

Furthermore, internal symmetry is important in understanding the stoichiometry of protein assemblies. Uneven stoichiometry assemblies are those with an unbalanced number of each entity type in the complex and occur rarely in the biological environment. It was previously reported that up to 40% of all protein assemblies with uneven stoichiometry in the PDB can be explained by the presence of internal symmetry in one or multiple of the subunits in the complex [41]. One such example is the artificial complex of Bowman-Birk inhibitor from snail medic seeds with bovine trypsin, which has an A2B stoichiometry (Fig 5). Although the complex is asymmetric, considering the internal symmetry of the inhibitor shows that the assembly is structurally comparable to an even A2B2 assembly with C2 overall symmetry. This property has also functional consequences, since the binding of two trypsin proteins symmetrically allows the inhibitor to efficiently induce dimerization and block the peptidase activity. Symmetry is characteristic of biological assemblies and can be considered by methods, like EPPIC, in order to predict the biological assembly in the context of crystal latices [42]. Including internal symmetry in these methods could further improve predictions for some known cases like, for example, uneven stoichiometries.

**Fig. 5.**
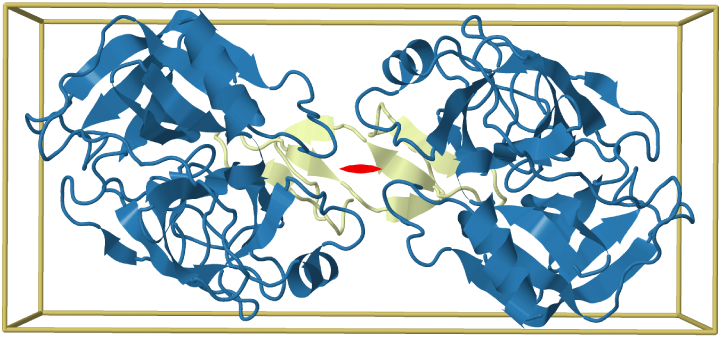
Uneven stoichiometry as a consequence of internal symmetry. The Bowman-Birk inhibitor from snail medic seeds (yellow) forms a complex with two bovine trypsin subunits (blue) in an uneven (A2B) stoichiometry and asymmetric assembly (2ILN). However, a 2-fold symmetry axis (red) can be identified in the complex when the internal symmetry of the inhibitor is taken into account, showing that the complex is equivalent to one with even (A2B2) stoichiometry.

### Open Symmetry

The majority of proteins have closed symmetry. In the case of quaternary structures, this is expected since homooligomers with open symmetry are disfavored due to their aggregation potential [43]. However, this is not the case for internal symmetry due to the ability for terminal repeats to diverge and avoid undesirable homotypic interactions. The most general formulation of open repeats in the literature is that of superhelical symmetry, where the repeating unit is simultaneously translated along a helical path (curvature) and rotated around this path (twist) [27]. `CE-Symm` cannot identify superhelical symmetries, where both curvature and twist are relevant, because of thefundamental limit of the method to find a single symmetry axis (or multiple independent axes). However, we observe that the majority of structures containing tandem repeats that are classified as superhelical in the literature (solenoids) can be approximated with a single axis of symmetry by `CE-Symm`. They fall in one of the following four conditions: i) the twist is negligible (relative to `CE-Symm` tolerances); ii) the curvature is negligible; iii) both twist and curvature are negligible; or iv) the twist is much larger than the curvature. In all those cases, `CE-Symm` can identify the symmetry in the structures and annotate them as helical, translational or open rotational symmetries.

For instance, from the 18 solenoid protein representatives from table 1 in Kobe and Kajava [27], in 10 either the twist or the curvature are reported to be small (helical symmetry applies), in 5 both the twist and curvature are annotated as small (translational symmetry applies), and the remaining 3 structure representatives have irregular twist (asymmetric applies). Although many folds are classified as superhelical, only a small number have regular repeats but do not fit into one of the above categories. Therefore, in practice `CE-Symm` can also be a good method for identifying, classifying and characterizing solenoid and other repeat proteins with open symmetry. We hypothesize that the low prevalence of actual superhelical symmetry in proteins could be a consequence of the benefit in conserving interfaces between adjacent repeats.

## Conclusion

We have extended our internal symmetry analysis tool in order to improve its usability, capabilities and the interpretability of results. In addition to detecting symmetry in protein structures, the tool can identify corresponding residues of the protein from each repeating element and the symmetry operations between them. `CE-Symm` 2.0 adds broad capabilities for the detection of all types of internal symmetry, providing information about the type and order of symmetry and the repeat boundaries. The alignments between the repeats are eminently useful in identifying conserved and differential features between repeats, and can be applied to understanding protein function and evolution.

The ability to run `CE-Symm` recursively to detect multiple axes of symmetry allows both higher-order point group symmetries to be identified and non-point group hierarchical symmetries. The simultaneous visualization of this rich information can lead to a better understanding of the structure and provide information about multiple duplication events. One limitation of `CE-Symm` 2.0 is that it does not yet integrate quaternary symmetry detection into it’s hierarchy. While it is possible to run the program on biological assemblies, it will have poor performance and may mistakenly fail to detect symmetry. Rather, methods specific to quaternary symmetry detection should be integrated with CE-Symm to provide this feature.

`CE-Symm` has been optimized for finding structures with global symmetry. While it does search for insertions, the length dependence of TM-Score means that structures with large insertions or multi-domain queries may not meet the default score thresholds. For multidomain proteins it may be needed to perform domain decomposition prior to running `CE-Symm`. Since it is based on CE rigid body alignment, the tool is also unlikely to detect all repeats in structures with conformational changes in some repeats, or with non-sequential rearrangements like circular permutations between repeats. Another limitation is the requirement for a consistent orientation between equivalent repeats. While for some applications preserving a conserved interface between repeats is desirable, there are many cases with large and functionally significant changes in repeat orientation (e.g. in many solenoid proteins).

Determining whether the high prevalence of internal symmetry in protein structures is predominantly a consequence of thermodynamic selection or an indication of the history of protein evolution remains an open question. Here, we have presented examples where internal symmetry is a result of evolution and tied to functional consequences, and how our tool can help researchers in the protein evolution, classification and annotation fields. The `CE-Symm` source code has been integrated into the BioJava library [44] and is freely available on GitHub. In the future, we would like to integrate `CE-Symm` into leading bioinformatics resources for protein analysis.

## Author’s contributions

P.B. initiated the CE-Symm project and provided direction. S.B., A.L, P.R. and A.P. developed code for the CE-Symm tool. S.B. and A.L. designed, implemented and benchmarked the algorithmic extensions of the method and performed the analyses presented in this article. S.B., A.L. and G.C. wrote the manuscript. All authors reviewed and approved the final version of the manuscript.

## Acknowledgements

We dedicate this article to Guido, catalyst and indispensable part of this project, who sadly left us before its completion.

We thank Philippe Youkharibache for testing and suggesting new features for `CE-Symm` and Jose Duarte for useful discussions during the development of the method. Layla Hirsh Martinez kindly provided RepeatsDB-lite results on our benchmark and contributed helpful insights to the research. We also thank the developers that contributed to the BioJava library, which has been fundamental throughout the development of `CE-Symm`.

This research was supported in part by the Intramural Research Program of the National Center for Biotechnology Information, National Library of Medicine, National Institutes of Health (support to S.B and P.B), and by the National Science Foundation, National Institutes of Health, and U.S. Department of Energy [DBI-1338415] (support to A.P., P.R., and P.B.). Financial support to G.C. from the Swiss National Science Foundation and the Research Committee of the Paul Scherrer Institute is gratefully acknowledged. Additional support for S.B from COST Action BM1405 and COST Switzerland SEFRI project IZCNZ0-174836.

## Supporting information

**S1 Fig.**
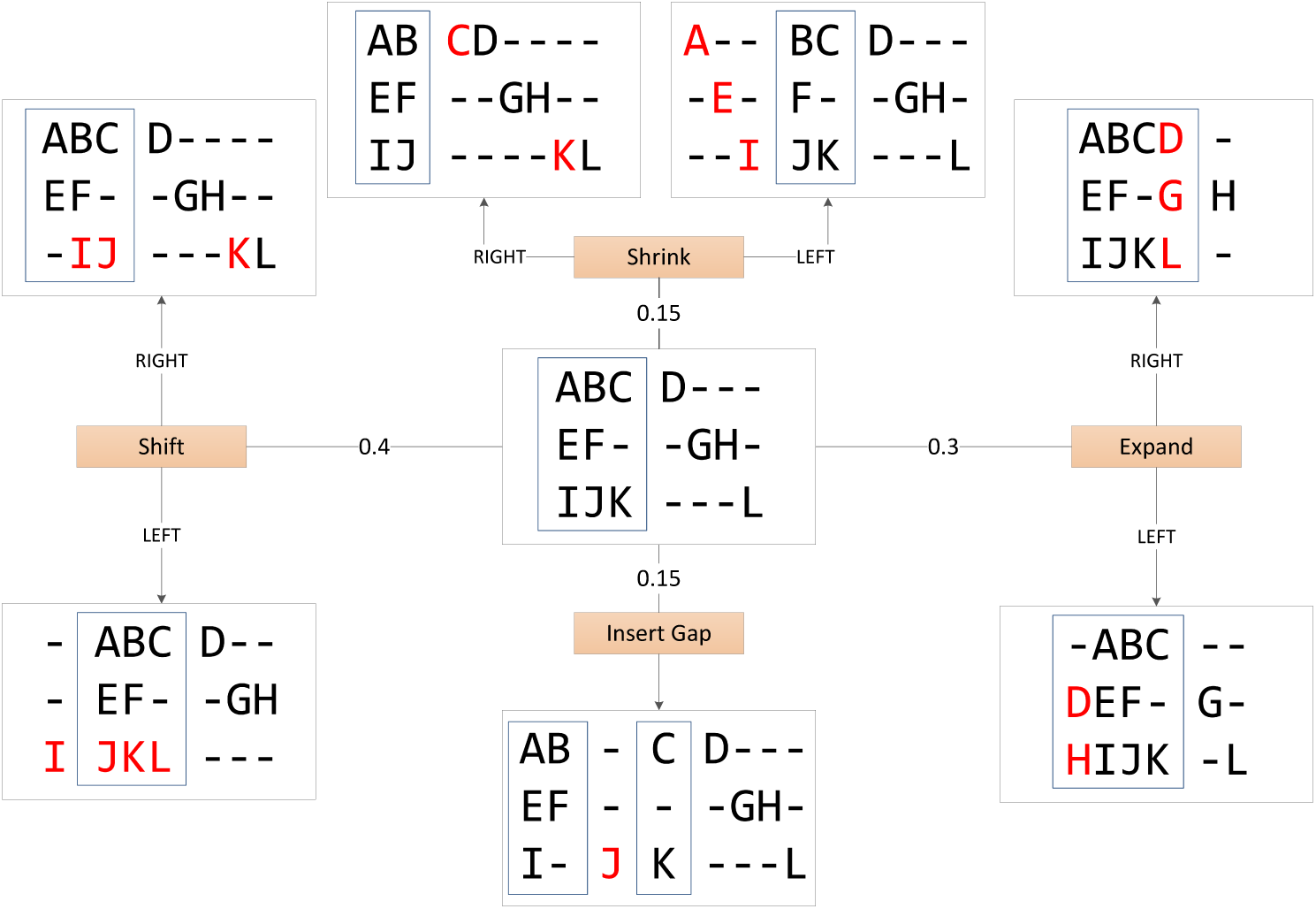
Schematic representation of the Monte Carlo optimization moves. The starting alignment is shown in the center. The probability of each of the moves are indicated along the edges.

**S2 Fig.**
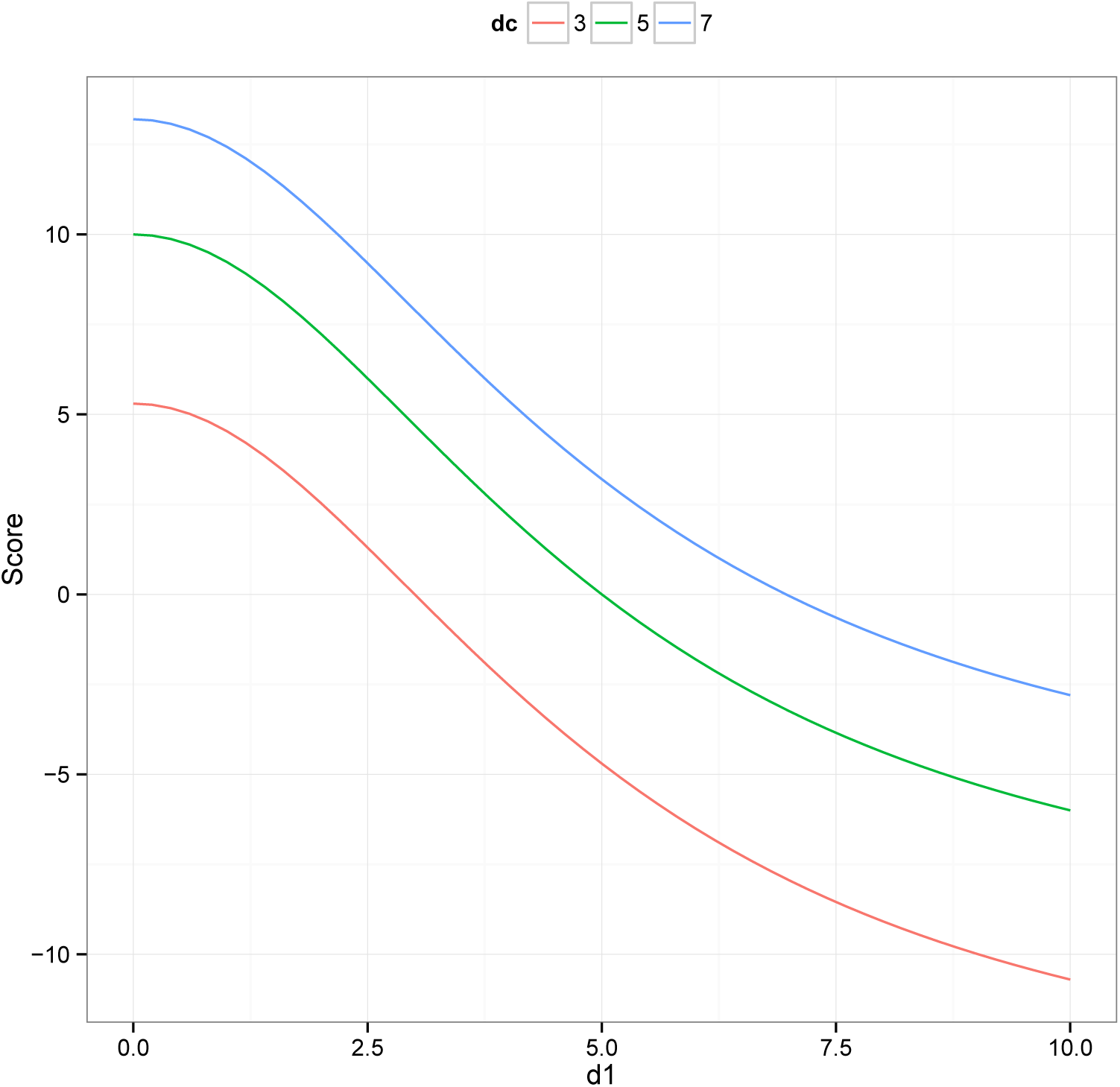
Score function for the Monte Carlo optimization procedure.

**S3 Fig.**
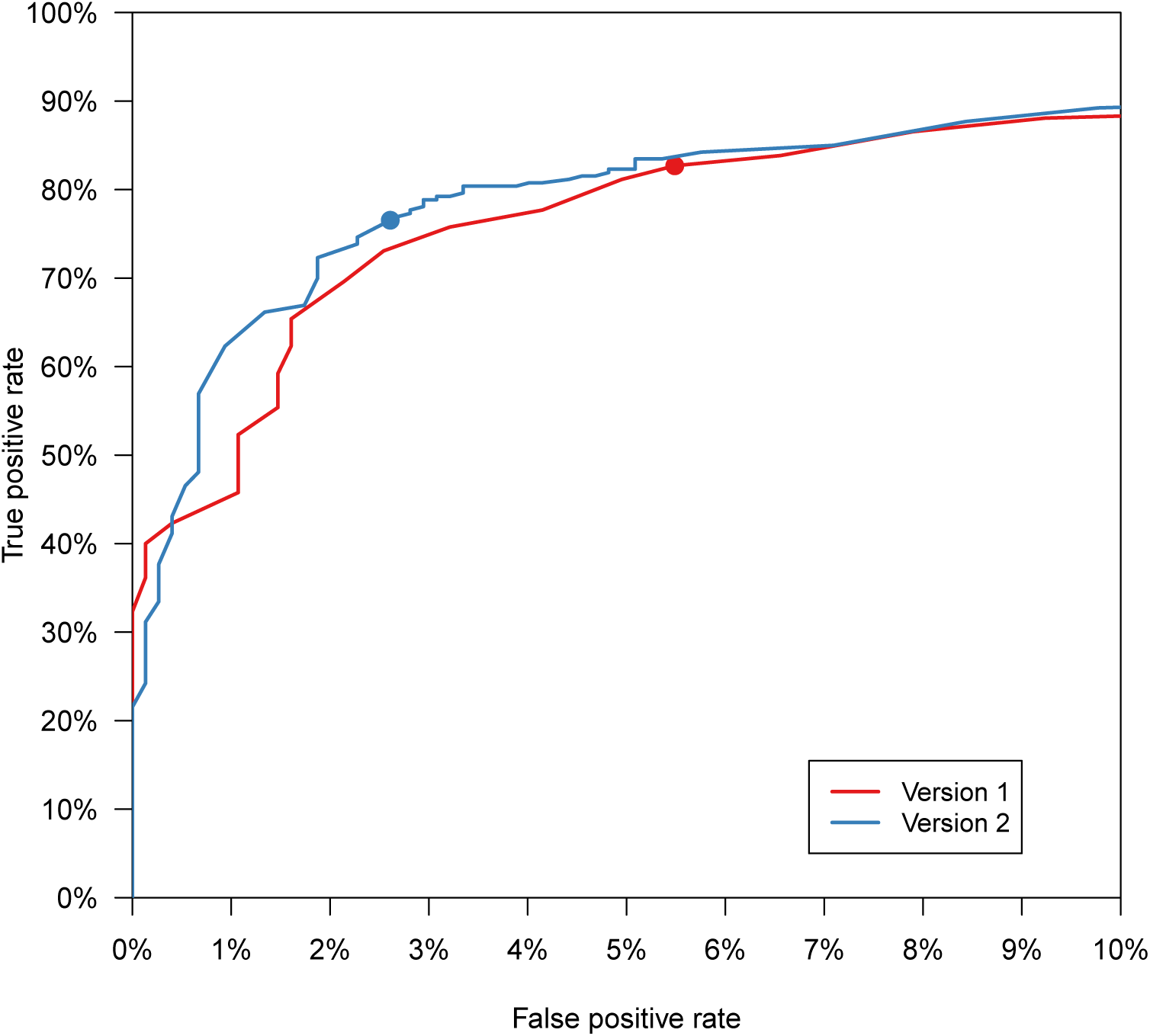
Comparison of the ROC curves of the symmetry detection for the old (Version 1) and new (Version 2) versions of CE-Symm. Differences in the ROC curves are not significant. The dots indicate the sensitivity and specificity at the default TM-score threshold (0.4).

**S4 Fig.**
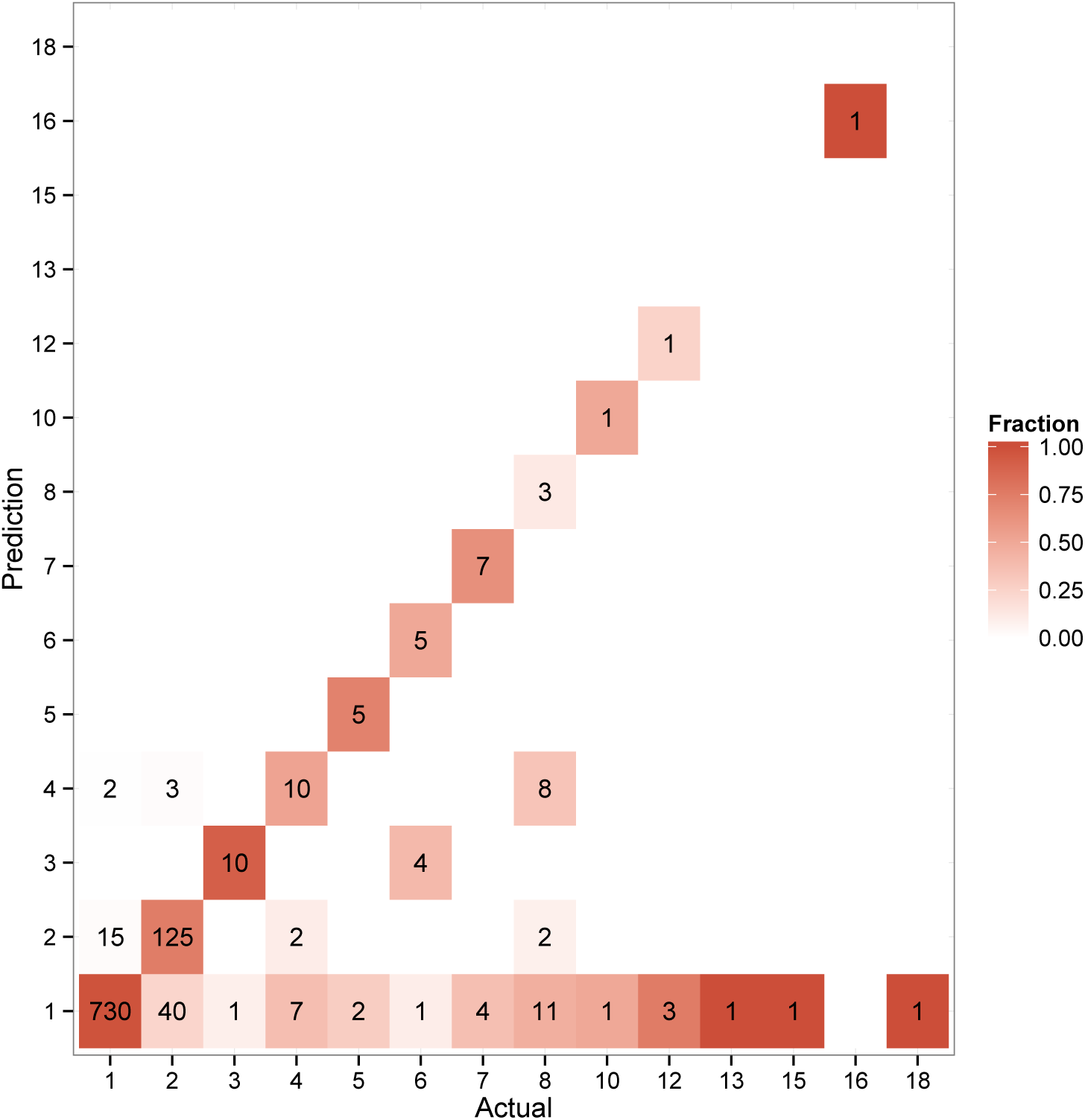
Confusion matrix of actual and predicted CE-Symm symmetry orders of the structures in the benchmark. Entries of the matrix are colored by the recall of each symmetry order (columns).

**S5 Fig.**
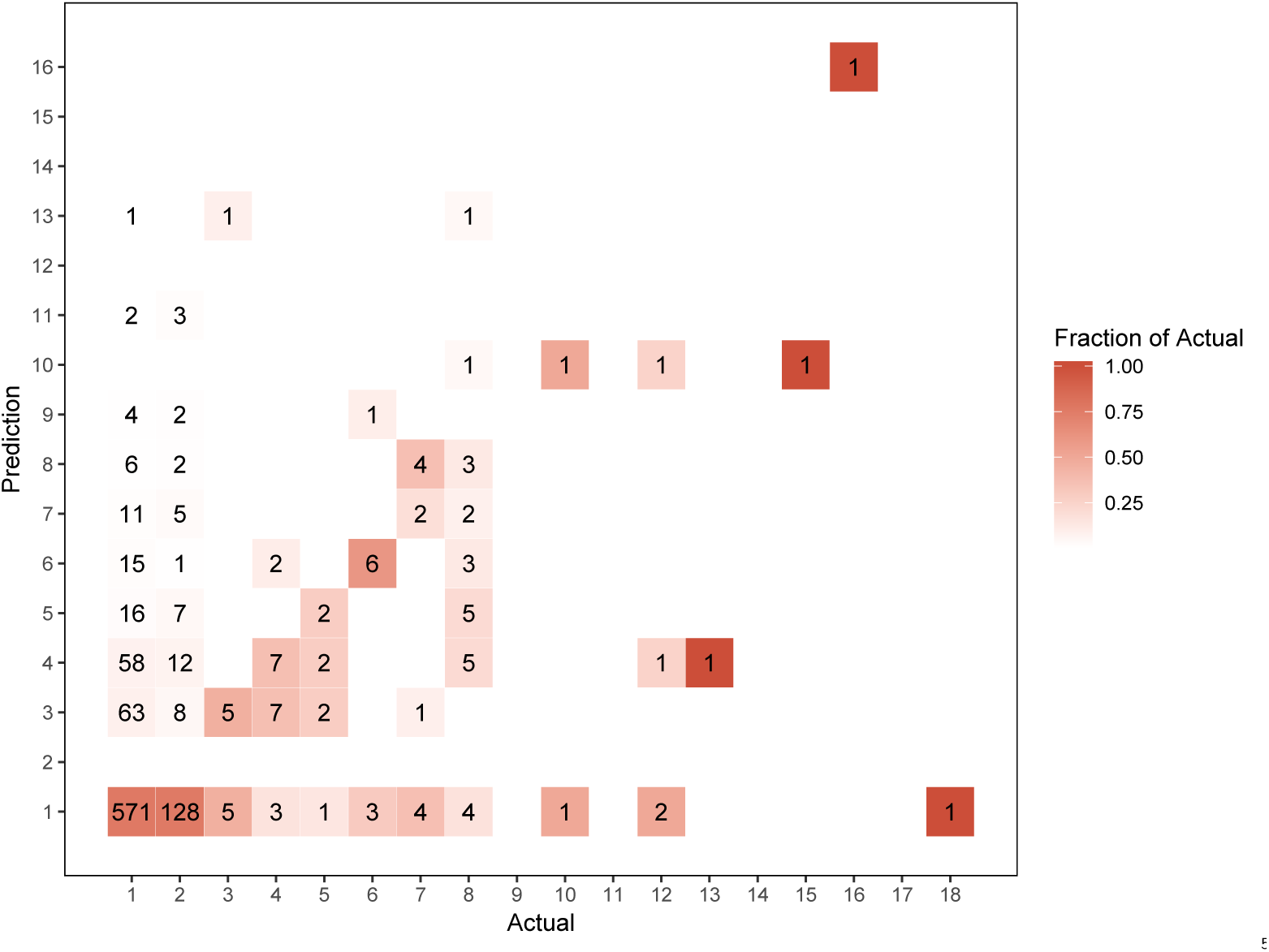
Confusion matrix of actual and predicted RepeatsDB-lite symmetry orders of the structures in the benchmark. Entries of the matrix are colored by the recall of each symmetry order (columns).

**S1 Tab.** **Summary of the updated annotations in the internal symmetry benchmarking dataset**.

| Type | Count | Percentage |
| --- | --- | --- |
| **Asymmetric** | **747** | **74.2%** |
| **Rotational** | **214** | **21.2%** |
| C2 | 160 | 74.8% |
| C3 | 10 | 4.7% |
| C4 | 2 | 0.9% |
| C5 | 3 | 1.4% |
| C6 | 9 | 4.2% |
| C7 | 10 | 4.7% |
| C8 | 20 | 9.3% |
| **Dihedral** | **18** | **1.8%** |
| D2 | 14 | 77.8% |
| D3 | 1 | 5.6% |
| D4 | 2 | 11.1% |
| D5 | 1 | 5.6% |
| **Helical** | **11** | **1.1%** |
| H2 | 9 | 81.8% |
| H3 | 2 | 18.2% |
| H10 | 1 | 9.1% |
| **Superhelical** | **2** | **0.2%** |
| **Repeats** | **15** | **1.5%** |

**S2 Tab.Summary of results from running `CE-Symm` and RepeatsDB-lite on the 1007 SCOP domains of the benchmark.**Tab-delimited file giving the symmetry and number of repeats found for each domain. Also available at https://raw.githubusercontent.com/rcsb/symmetry-benchmark/master/domain_symm_benchmark/domain_symm_benchmark_results.tsv.

**S3 Tab.** Results on RepeatsDB reviewed entries. Tab-delimited file giving the 3495 entries, with annotations about the number of domains and the `CE-Symm` 2.0 results. Also available athttps://raw.githubusercontent.com/rcsb/symmetry-benchmark/master/repeatsdb-lite/repeatsdb-benchmark.tsv.

| Method | Precision | Cramer V |
| --- | --- | --- |
| GRAPHCOMPONENT | 0.598 | 0.652 |
| DELTAPOSITION | 0.783 | 0.728 |
| ROTATIONANGLE | 0.754 | 0.642 |

**S4 Tab.Performance measures of the symmetry order detection methods for domains in the benchmark dataset with closed symmetry.**Precision measures the total fraction of correct predictions and Cramer V measures the correlation between actual and predicted classes. Both measures have values in the [0,1] interval, where 1 means perfect precision and correlation.

---

**S1 Text. Supplementary Methods.** PDF file with Supplementary Methods.

## Supplemental Methods

### Order detection and refinement methods

The Rotationangle is a geometric method for determining the order in cases of closed symmetry. It is based on the angle of rotation, which can be calculated from the superposition operator (see Additional file 6 of [20]). The distance between a measured angle of rotation, *θ*, and the closest theoretical angle of rotation for order *k* is given by a triangle wave of frequency *k*:

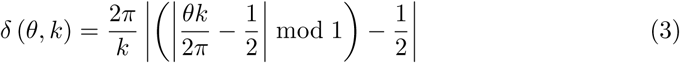

Note that the equation given in [21] (there notated *ɛ*(*θ*)) assumed that the ideal rotation would be 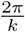 and neglected to account for other multiples of this. The triangle wave equation above properly accounts for the multiple possible ideal angles in rotationally symmetric structures.

The best-fit order is then the *k* that minimizes this distance, up to some maximum order.

Both the GraphComponent and DeltaPosition methods can be understood as operations on a directed graph, where the set of residues form nodes and are connected by an edge if the alignment aligns one onto the other. Order detection is performed on the initial graph, while refinement consists of modifications of the graph. The initial self-alignment has few restrictions other than all nodes having out-degree of at most one; the presence of a circular permutation in the self-alignment permits several residues to align to a single target residue. Refinement is complete when the remaining nodes consist of either linear paths (open symmetry) or simple cycles (closed symmetry), each with *k* nodes. These are then sorted according to the protein sequence and converted into columns of the output multiple alignment.

For the GraphComponent method, the order is first determined as the most frequent size of connected component. In the refinement step, the graph is then modified by discarding all nodes not belonging to a path of *k* nodes. The largest subset of the remaining paths is chosen such that the sequence order of the protein is preserved in the multiple alignment. This is done by checking whether a pair of paths are “compatible”, meaning they can be sorted *a* < *b* such that

*a1* < *b*_1_ < *a2* < *b*_2_ < … < *a*_*k*_ < *b*_*k*_. The connected component which is compatible with the most other components is selected greedily for inclusion in the refined multiple alignment. While this procedure reduces the alignment length, it was found to usually leave sufficient columns to seed the optimization step.

The DeltaPosition method attempts to better handle difficult cases with closed symmetry, where errors in the self-alignment can lead to the alignment graph becoming highly connected. For each node *x*, let *f* ^*k*^(*x*) denote the node reached by following the path from *x* through *k* nodes. In a good alignment with the correct order *k*, many paths will form cycles of *k* nodes, so *f* ^*k*^(*x*) = *x*. In noisier alignments, *f* ^*k*^(*x*) will not close a cycle, but will still be close to *x* in terms of the sequence position. Thus, the position distance Δ(*x*) = |*x* — *f* ^*k*^(*x*)| measures how close a particular residue is to forming a cycle. To determine the order, *k* is chosen up to a maximum order (8 by default) so that it minimizes

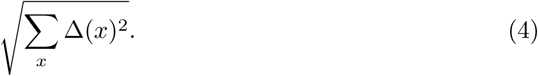

After detecting the order, the DeltaPosition refines the alignment until it consists only of cycles of *k* nodes. In each step, a linear path of *k* nodes is selected from the graph to become a closed cycle. The path is chosen greedily according to the Δ(*x*) of it’s start node, and the outgoing edge from the *k*th node is modified to point back to *x*. During every step, only cycles are considered which are compatible with previously selected cycles with respect to the protein sequence order. Finally, any nodes not belonging to cycles are discarded and the refined multiple alignment corresponding to the set of cycles is output.

Several additional order detection variants were considered during the development of `CE-Symm` 2.0 but discarded after analyzing their performance [45].

### RepeatDB-lite comparison

RepeatDB-lite was run on the 1007 domains of the benchmark. Five domains produced errors since they are low resolution structures with only CA atoms positioned. In all cases, higher resolution structures of the same proteins are available which include all atoms. Thus, the following five cases were substituted into the benchmark. (`CE-Symm` results are the same on both low and high resolution structures for all cases.)

**Tabel.1.**

| Benchmark Domain | Replacement Structure |
| --- | --- |
| d1i95b_ | d2uubb1 |
| d1i96v_ | 5LMN.X:83-169 |
| d2rdo81 | d1isea_ |
| d2rdow1 | d2gycu1 |
| d3b5zd2 | d3b60d2 |

All substitutions have identical sequence and have very similar structures. SCOPe 2.01 domains were used where possible. The exception to this is `d1i96v_`, of which no other structures had been classified by SCOPe.

